# Long-term cellular cargo tracking reveals intricate trafficking through active cytoskeletal networks

**DOI:** 10.1101/2022.03.23.485568

**Authors:** Jin-Sung Park, Il-Buem Lee, Hyeon-Min Moon, Seok-Cheol Hong, Minhaeng Cho

## Abstract

A eukaryotic cell is a microscopic world within which efficient material transport is essential. To examine cargo transport in a crowded cellular environment, we tracked unlabeled cargos in directional motion in a massively parallel fashion using the interferometric scattering microscopy. Our label-free, cargo-tracing method revealed not only dynamic cargo transportation but also the fine architecture of the actively used cytoskeletal highways and the long-term evolution of the associated traffic at sub-diffraction resolution via a myriad of molecular strokes. Cargos experience a traffic jam, but they have an effective strategy to circumvent it: moving together in tandem or migrating collectively. All taken together, a cell is an incredibly complex and busy space where the principle and practice in transportation intriguingly parallel those of our macroscopic world.

---

Intracellular cargo transport plays a critical role in maintaining the essential functions of living cells^1,2^. For cargo transport, vesicles are constantly formed as a container at the cell membrane, endoplasmic reticulum (ER), and Golgi apparatus^3,4^ and transported by motor protein-driven activity on cytoskeletal highways^5,6^. During this transport, cargos exhibit rich and intricate dynamic events such as directional movement, intermittent pausing, turn-around, and road change at the intersection of cytoskeletal highways^7,8^, paralleling manufactured vehicles operating in the urban road network (Fig. 1a). It is, however, still unclear how these cargos overcome general traffic problems such as traffic congestion, possibly severer in the heavily crowded cellular environment^9,10^.

**Fig. 1.**
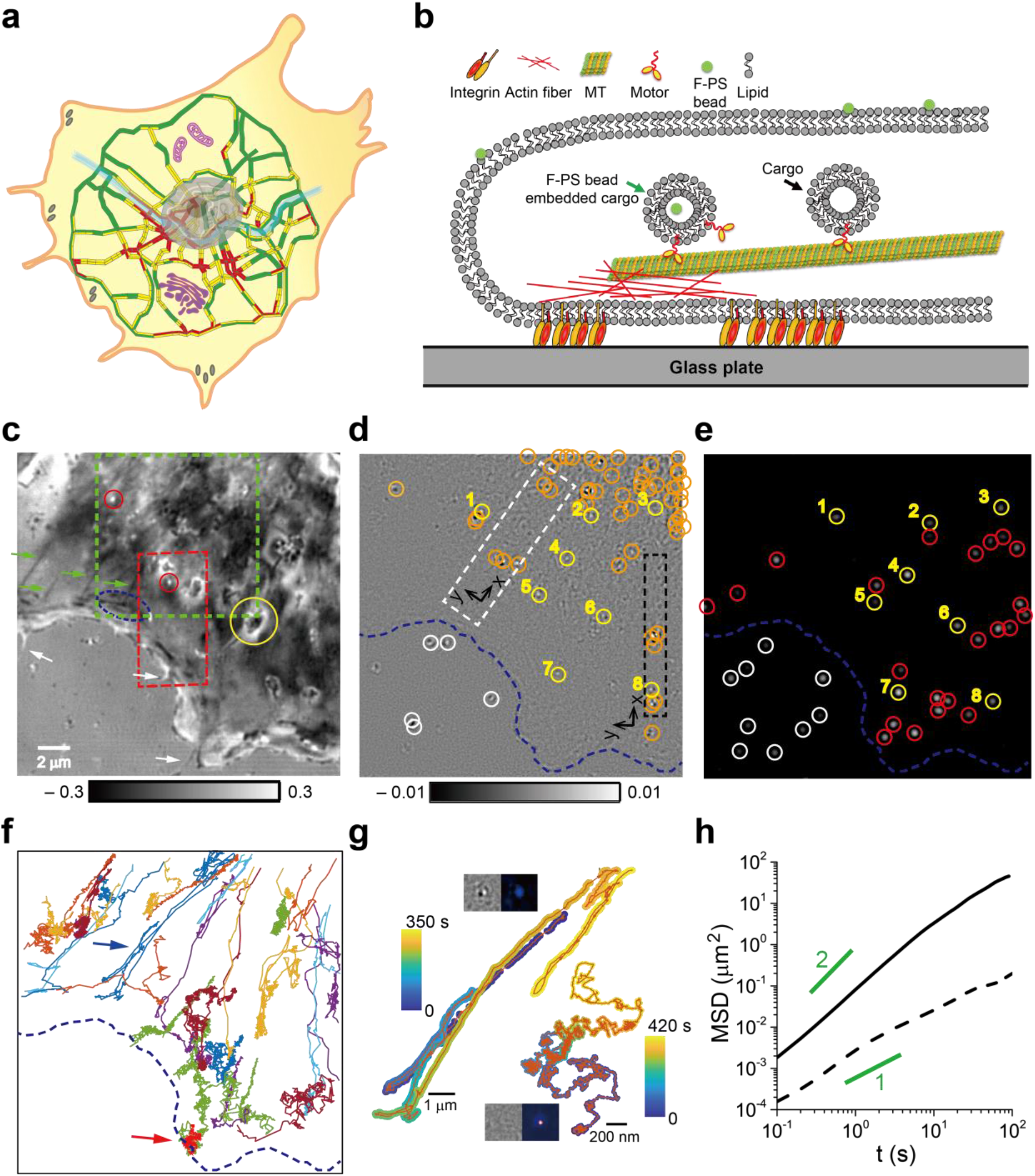
Intracellular cargo transport in a complex cellular environment. **a,** Schematic illustration of intracellular trafficking based on the traffic road map of the metropolitan region of Seoul, Korea. The colors on each road section indicate the degree of traffic congestion with red, yellow, and green indicating heavy, medium, and light traffic, respectively. The large gray oval in the center, complex purple structures sideway, and gray spots at the cell boundary represent the nucleus, cytoplasmic organelles (mitochondria and Golgi apparatus here), and focal adhesions, respectively. **b**, Schematic picture for intracellular cargo transport along a cytoskeletal filament. Cargos bearing a f-PS bead can be detected simultaneously with iSCAT and fluorescence microscopies. **c**, SBR-iSCAT image showing various intracellular structures. Here, the filamentous structures (green arrows) and their intersection (blue oval) are shown in dark contrast. The large and small globular structures (yellow and red circles, respectively) are visible in dark and bright contrast, respectively. Filopodia (white arrows) are also observed at the cell edge. **d**, TD-iSCAT image of the same area as in (c) to show the locations of moving cargos. **e**, Fluorescence image of the same area to show the locations of f-PS beads inside and outside the cell. Yellow circles marked with the same number in (d) and (e) indicate co-localized cargos and f-PS beads, respectively, signifying f-PS bead bearing cargos. Orange and red circles mark spots observed in one channel only (orange in (d) and red in (e): cargos and f-PS beads, respectively). White circles in (d) and (e) show spots outside the cell. **f,** Trajectories of 32 different f-PS beads drawn in different colors, tracked at 10 Hz by fluorescence microscopy. **g,** Trajectories of two f-PS beads that represent (bi-)directional or Brownian motions, indicated by blue and red arrows in (f), respectively. Insets: TD-iSCAT image (left) and fluorescence image (right) of each tracked particle; directional and Brownian particles are shown in the upper and lower insets, respectively. **h,** MSD-vs.-time curves of the particles shown in (g). Solid and dashed lines represent the MSD curves by directional and Brownian particles with the diffusion exponent of α ~ 1 and 1.5, respectively. Lines with α = 1 and 2 are drawn in green for visual guidance. In (d-f), the cell boundary was shown in dashed line (blue) for visual guidance. See also Extended Data Fig. 2, Supplementary Video 1 and 2.

Our current understanding of cell metabolism has been brought about by fluorescence-based imaging techniques. The specificity, sensitivity, multiplicity, and super-resolution capability of fluorescence microscopy are critical assets as a powerful tool for cell study. Various issues about cargo transport have been addressed, including motor protein dynamics in cytoplasmic environments^11,12^, vesicle transport in 3D^8^, cargo dynamics at cytoskeletal junctions^7,13^, interactions of cytoskeletal network and organelles in transport at high spatio-temporal resolutions^14,15^. Fluorescence-based methods are, however, fundamentally limited in long-term live-cell imaging due to photobleaching. Besides, they leave unlabeled cellular constituents invisible like dark matter, missing all the information about the local environment except labeled target molecules, although the unlabeled majority likely dictates the apparent behavior of target molecules.

Here, we report the universal feature of intracellular traffic flow, revealed by simultaneously tracking many unlabeled cargos transported along cytoskeletal highways in a highly parallel fashion and for an indefinitely long time using the interferometric scattering (iSCAT) microscopy. The label-free and high-speed imaging by iSCAT permits us to capture the quantitative and physical details about cargo transport and to acquire a realistic picture of nanoscale logistics within living cells. The enormous amount of cargo localization data (> 10^8^) enables us to reconstruct the fine architecture of cytoskeletal meshwork and visualize the temporal evolution in traffic flow along the active cytoskeletal highways. To our amazement, cells intrinsically have an efficient transport strategy to avoid an intracellular traffic jam by forming a train of cargos or collectively moving in the same direction, closely resembling our daily life.

## Cargos in directional motion are tracked in a complex cellular environment via iSCAT microscopy

As a notable label-free imaging technique, the iSCAT microscopy has drawn much attention due to high detection sensitivity^16–19^, ultrafast tracking of dynamic nanoparticles^20,21^, 3D imaging capability^21–24^, and biological applications towards live-cell imaging^25–28^. To identify cellular vesicles and learn how they appear in our home-built iSCAT microscopy (Extended Data Fig. 1), we fed COS-7 cells with 20-nm fluorescent polystyrene (f-PS) beads and visualized them with fluorescence and iSCAT microscopies simultaneously (Fig. 1b, d, e, g).

In the static background-removed iSCAT (SBR-iSCAT) image (Extended Data Fig. 2a, c), various subcellular structures are visible with enhanced contrast (Fig. 1c). Besides filamentous structures, large and small globular objects are easily identifiable (yellow and red circles, respectively; Fig. 1c). Filopodia are also clearly shown at the cell edge (white arrows; Fig 1c). Nanoscale cargos may still be challenging to detect in the presence of complicated cytoplasmic background. To separately detect the scattering signals of nanoscale cargos in the cytoplasmic background, we should resort to the dynamic nature of cargos. One remarkable feature in cellular cargo transport is (bi-)directionality with intermittent pauses, as in the case of motor vehicles in busy two-way roads. Other cytoplasmic constituents appear relatively static or randomly jiggling. To discern mobile cargos from such static objects and sort out directional cargos from random walkers, we employed the time-differential iSCAT (TD-iSCAT) method (Fig 1d, Extended Data Fig. 2b, d).

Among f-PS beads identified by fluorescence microscopy (Fig. 1e), only eight of them are co-localized in the TD-iSCAT image (Fig. 1d). The trajectories of 32 f-PS beads were acquired by tracking their fluorescent signals at 10 Hz (Fig. 1f). These trajectories could be divided into two distinct types, directional or Brownian. From the representative data in each type (Fig. 1g, Supplementary Video 1, 2), the diffusion exponents (α) that are the slopes of the MSD(mean squared displacement)-*vs*.-time graphs are approximately 1.5 and 1 for directional and Brownian motions, respectively, reflecting their markedly different dynamic characteristics (Fig.1h). Interestingly, we found that TD-iSCAT imaging only captures f-PS beads in directional motion: the insets (Fig. 1g) show that in iSCAT image, a Brownian f-PS bead, sharply imaged in the fluorescence channel, is obscure while a directional f-PS bead with dimmer fluorescence is clearly detected, as shown in the lower and upper insets, respectively. Thus, intracellular cargos are likely the main target of TD-iSCAT imaging as many native non-fluorescent cargos that lack f-PS beads are readily visualized by TD-iSCAT as indicated by orange circles (Fig. 1d).

## Cargo-localization iSCAT microscopy reveals the underlying active cytoskeletal highways

As described above, iSCAT directly captures the dynamic feature of individual cargos transported along cytoskeletal filaments without external labels. As a high-speed and time-unlimited label-free imaging technique, iSCAT produces a large amount of imaging data and thus provides a unique opportunity for statistical analysis. We note that cargos in directional motion are confined on cytoskeletal tracks and the cargo location is a faithful proxy for a cytoskeletal foothold with an error of cargo size. In order to calculate the localization precision for single cargos revealed in TD-iSCAT images, the pair of bright and dark spots that represents a single cargo was fitted with a pair of two 2D Gaussian functions (Extended Data Fig. 3). For bright and dark spots, the value of signal-to-noise ratio (SNR), 6 and 4, allows localization of the center of their Gaussian peaks to be within 10 and 15 nm in precision, respectively. Thus, by collecting a vast number of cargo locations, one can reconstruct the underlying cytoskeletal network. We define cargo location by the center of the bright spot (Extended Data Fig. 4a) using the MOSAIC ImageJ plugin^29^. We could visualize their underlying cytoskeletal tracks by accumulating a vast number of localization points (~100,000) obtained from 50,000 sequential images (Extended Data Fig. 4b). Similarly, the cytoskeletal network in lamellar and lamellipodium in a COS-7 cell was reconstructed from ~ 10-million localization points obtained from ~ 90,000 consecutive frames taken at 50 Hz (Fig. 2a).

**Fig. 2.**
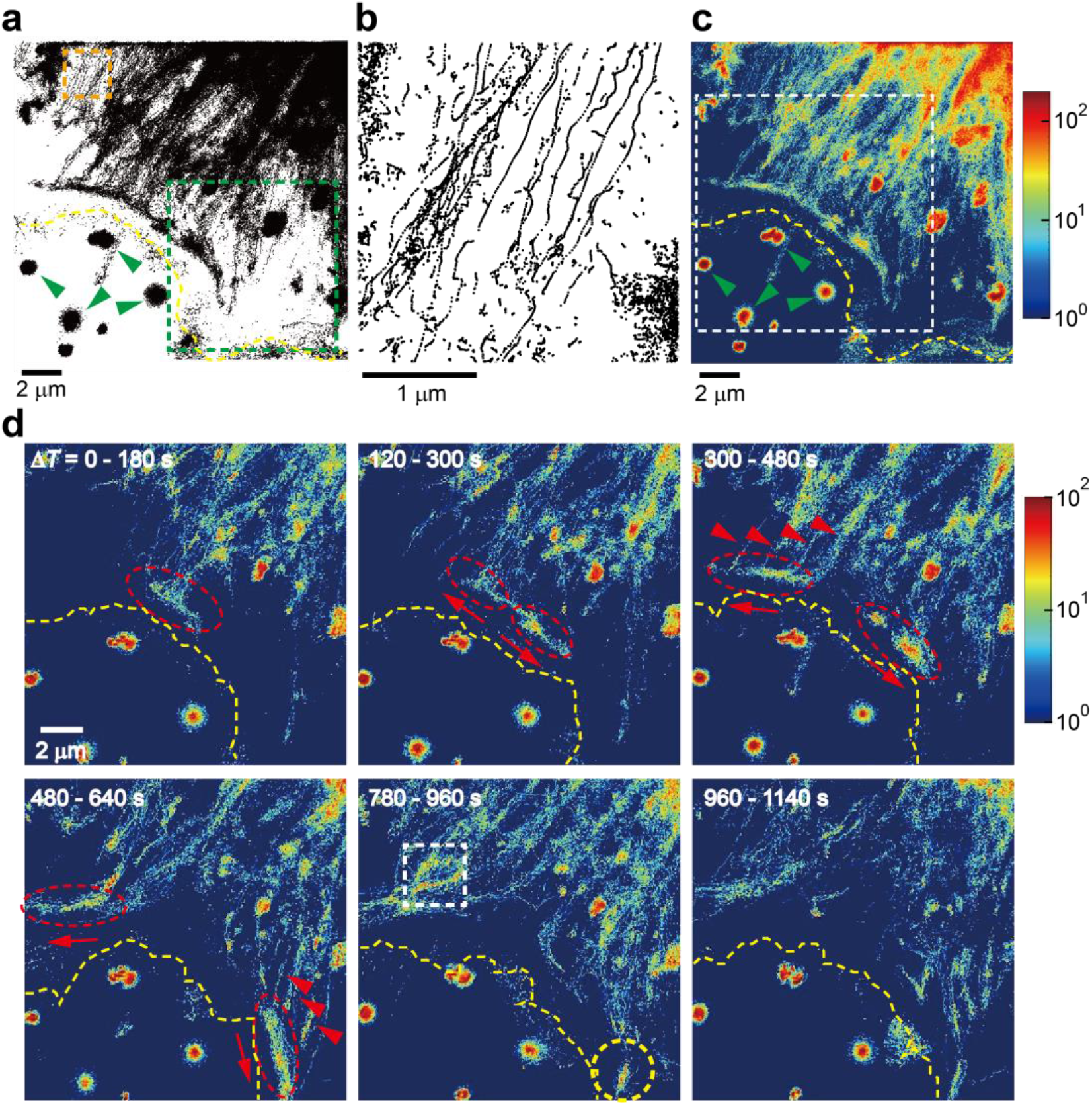
Temporal evolution of cargo flow on active cytoskeletal highways. **a,** Cytoskeletal network reconstituted by cargo localization performed in TD-iSCAT images (Fig. 1d). About ten-million cargo positions from ~ 90,000 consecutive frames taken at 50 Hz *(ΔT* ~ 30 min) were used to make this map. **b,** Zoomed-in image of the orange-boxed area in (a), showing the fine structure of cytoskeletal highways. **c,** Traffic density map. Color bar indicates the total numbers of cargos detected at each pixel for ~ 90,000 consecutive frames. **d,** Sequence of cargo-localization density maps integrated for 9,000 consecutive frames (*Δt* = 180 s) taken from the white-boxed region in (c). Only six characteristic images are presented here. The packet of cargos (red oval in the 1^st^ image) was branched out at the intersection of cytoskeleton highways. Then, these two packets rushed away in the opposite directions to the protruding areas (1^st^ to 4^th^ image). During this process, the traffic density increases as other packets are combined at the intersection of cytoskeleton highways (red arrow heads in the 3^rd^ and 4^th^ images). Once a packet reached the protruding area, it was diminished (yellow circle in the 5^th^ image). In (a, c), green arrowheads indicate debris fluctuating outside the cell. In (a), (c), and (d), the cell boundary was shown in dashed line (yellow) for visual guidance. See also Extended Data Figs. 3-5, Supplementary Video 3.

The fine architecture of cytoskeletal networks was previously reported by fluorescence super-resolution microscopic imaging^7^. Such super-resolution images were obtained by point accumulation of fluorescent spots originating from photoswitchable fluorophores directly labeled to cytoskeletons, detecting the whole fluorophore-labeled cytoskeletal material. On the contrary, the cargo-localization iSCAT collects the positional information of individual mobile cargos, capturing the cytoskeletal ropeway selectively along which cargos actually traveled during the observation time (Fig. 2a,b). As expected, cytoskeleton highways are extended from the cell body (upper-right corner) to the cell boundary. These parallel highways are merged and spanned transversely along the cell boundary (Fig. 2a). Their long, straight shape implies that they are built of stiff microtubules with a large persistence length of 4 - 8 mm^30,31^. The direction of cargo movement is designated by the polarity of microtubule: the (+) and (-) ends of microtubule refer to the direction toward the cell body and cell boundary, respectively. The traffic density map (Fig. 2c) shows the total counts of cargo localization at each pixel through the whole recording of about 90,000 frames *(ΔT* ~ 30 min). As the ER, continuous membranous organelle surrounding the nucleus, is the departure station of vesicles carrying just synthesized proteins, the traffic density in the cell body is higher than in peripheral regions. The trajectory of an f-PS bead drawn by its fluorescence is overlaid by a white line drawn on the traffic density map and well-matched with the main traffic route identified in iSCAT imaging (Extended Data Fig. 4g).

We also analyzed the speed of cargos in directional motion along a cytoskeletal highway (Extended Data Fig. 4c, d). The TD-iSCAT frequently loses track of cargos whenever they intermittently stop (Extended Data Fig. 2d). Thus, it is a delicate task to obtain a long trajectory of cargo without the help of fluorescence imaging of the f-PS bead that co-propagates with the cargo. Here, we selected 9 and 10 cargos moving to the (+) and (−) end, respectively, continuously tracked over at least 3 s (> 150 consecutive frames) with iSCAT. Before evaluating the ‘instantaneous’ speed of cargo, we first acquired the ‘moving-averaged’ position to suppress noise in speed measurement. The x and y positions of cargo at 50 consecutive time points in a raw trajectory were averaged to yield the position of the cargo in the 50-point time-averaged (1 s) trajectory (Extended Data Fig. 5). Then, we calculated the instantaneous speed from the displacement made by the cargo for 0.02 s and collected such data to get the distributions of instantaneous speeds of cargos moving to the (+) and (-) ends (Extended Data Fig. 4e). From the distributions, the average speeds of cargos moving to the (+) and (-) end were 0.53 ± 0.28 μm/s and 0.45 ± 0.19 μm/s, respectively. Kinesin and dynein motors are the key players responsible for transporting cargos toward the (+) and (−) ends of a microtubule, respectively^32^. The speeds of those proteins have been measured using single-molecule techniques in vitro and in vivo, varying widely from 0.02 to 2 μm/s^33,34^, indicating that our estimates are within the range of measured values.

Next, we investigated the temporal evolution of the traffic of intracellular cargos, examining a sequence of traffic density maps, each acquired at the time interval of 60 s and integrated for *ΔT* = 180 s (Fig. 2d, Supplementary Video 3). At *ΔT* = 0 – 180 s, a packet of cargos was visible in the red-oval area, moving to the cell boundary. At the intersection of cytoskeletons, this packet was divided into two and driven in opposite directions along the cell boundary. During propagation, each packet was elongated by combining it with additional cargos approaching from the lamellar side, as indicated by red arrowheads in the 3^rd^ and 4^th^ images. Once it reached the extruding area at the edge of the lamellipodium, this traffic flow gradually disappeared in the 5^th^ and 6^th^ images. A growing filopodium viewed as a waterfront construction site, our observation implies that packets of cargos were funneled into the building site and terminated for unloading (Extended Data Fig. 6). More traffic packets were formed in yellow-boxed regions (Extended Data Fig. 6b). We found that the two vital traffic routes along the cell boundary appeared as thick dark bands (white arrows) in the SBR-iSCAT snapshot (Extended Data Fig. 6a), likely revealing the underlying structural element, actin arcs, for directed cargo movement.

## Cargos experience heavy traffic at the intersection of cytoskeletal highways

One of the advantages of iSCAT for live-cell imaging is its capability of providing the brightfield image of the entire area and simultaneously visualizing the backdrop as well as an event of interest. In our case, the behavior of mobile cargos can be better elucidated in the context of the local cellular environment, which is invisible in fluorescence imaging. iSCAT imaging reveals that a traffic jam occurs on the cytoskeletal highways in a live cell. We observed that a jammed intersection was temporarily formed (Fig. 3a) and many cargo-like spherical objects were found at this intersection in the zoomed-in SBR-iSCAT snapshot (left in Fig. 3a). Among them, the locations of 5 individual objects exactly matched those of moving cargos identified in the corresponding TD-iSCAT image (right in Fig. 3a). A few cargos in the TD-iSCAT image seemed to be attached together (red circle in Fig. 3a).

**Fig. 3.**
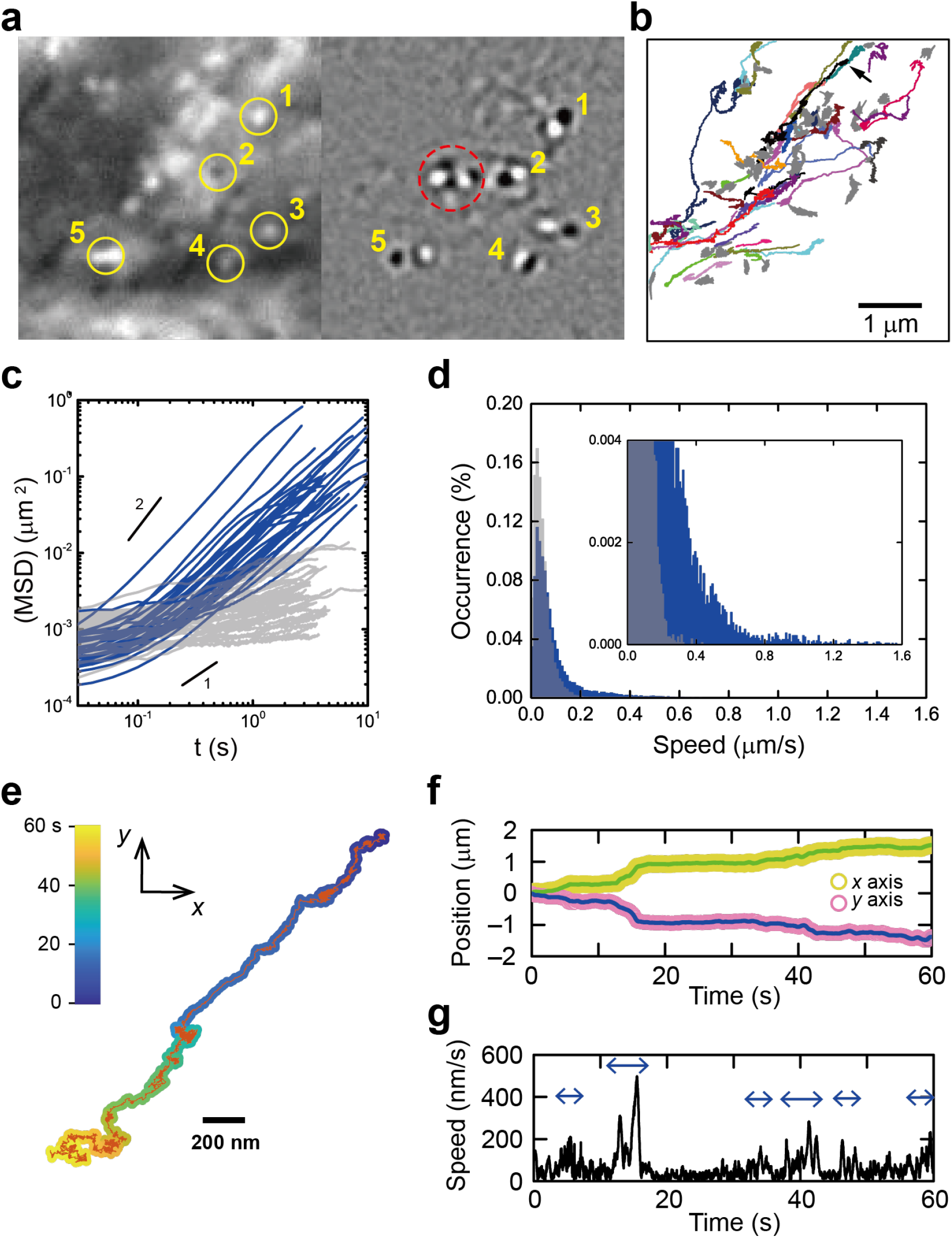
Traffic jam at the intersection of cytoskeleton. **a,** SBR-iSCAT (left) and TD-iSCAT (right) snapshot showing several adjacent cargos (boxed area drawn at t = 780 – 960 s in Fig. 2d). Both bright and dark spots in the SBR-iSCAT image (yellow circles marked by numbers) are associated with dynamic cargos shown in the TD-iSCAT image. In the red circle in the TD-iSCAT image, a few cargos are closely jammed. **b,** Total 83 trajectories of cargos continuously tracked over 500 frames (10 s) in 9,000 consecutive SBR-iSCAT images (180 s). Among them, the trajectories of 49 cargos with sub-diffusive motion were plotted in gray. The trajectories of 34 cargos showing directional motion were drawn in different colors. **c,** MSD-vs.-time curves of cargos traced in (b) (blue: cargos with directional motion, and gray: sub-diffusive cargos). **d,** Histogram of ‘instantaneous speed’ of cargos calculated from the two different groups, directional (blue) and jammed cargos (gray), in (b). The ‘instantaneous’ speed was calculated from time-averaged data points (averaged over 50 raw data points for Δt = 1 s) at the interval of 0.02 s for the raw trajectories drawn in (b). Directional cargos are dominant over jammed cargos in the regime of high-speed transportation (> 0.2 μm/s) (inset). **e,** A representative trajectory of a directional cargo (black trajectory indicated by the arrow in (b)) observed around the jamming area. **f, g,** Raw and time-averaged positions of cargo in (e) in x and y directions (x position: yellow (raw) and green (averaged); y position: purple (raw) and indigo (averaged)) (f) and its instantaneous speed (g). See also Extended Data Fig. 6, Supplementary Video 4.

Instead of tracking dynamic cargos from TD-iSCAT imaging, which is unusable for locally stalled cargos, we additionally tracked bright spots identified from SBR-iSCAT imaging to investigate the diffusion dynamics of cargos. Total 83 trajectories, continuously tracked over 500 consecutive frames (10 s) taken at 50 Hz, were overlaid (Fig. 3b). Among them, 34 cargos represented in different colors exhibited a directional movement with α ~ 1.5 as shown in the MSD-vs-time curve (Fig. 3b, c). Others were locally trapped and exhibited sub-diffusive dynamics with α < 1 (gray lines; Fig. 3b, c). The distribution of instantaneous speeds calculated from all trajectories of sub-diffusive cargos shows a positively skewed Gaussian shape with the mean value, 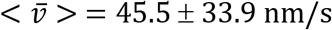 (Fig. 3d). Compared to that of locally trapped cargos, the measured average speed of directionally moving cargos is higher, with 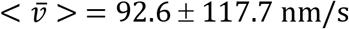, with a distribution shifted towards higher speed (Fig. 3d). One representative trajectory of a directionally moving cargo (Fig. 3e), however, showed that it was also congested intermittently throughout the observation time (Fig. 3f), and its instantaneous speed, except for short periods of time marked by arrows (Fig. 3g), is indistinguishable from those of sub-diffusive cargos.

## Dynamical phenomena revealed in cargo transport resemble commonplace events in transportation

The superb detection sensitivity of iSCAT allows various dynamic events to be revealed in the process of intracellular cargo transport. Interestingly, we found that some cargos move together as a pair, namely “dimeric cargo” (Fig. 4a). For a long observation time (> 130 s), cargos in such a dimeric form moved along the same path without being separated. However, its journey was not smooth due to many obstacles scattered around the path. The dimeric cargo moving toward the cell boundary could not move any further after colliding with a large cellular obstacle (time window ‘a’ in Fig. 4b). After stopping or randomly jiggling for a while, it changed the direction of motion and retraced its path (Supplementary Video 5). Then, its direction of motion changed once again by a head-on collision with another cargo moving oppositely along the same path (red arrow in Fig. 4b, Supplementary Video 6). The dimeric cargo stopped again at the surface of the large obstacle (time window ‘c’ in Fig. 4b), turned around, and then persistently propagated to the cell body (Supplementary Video 7).

**Fig. 4.**
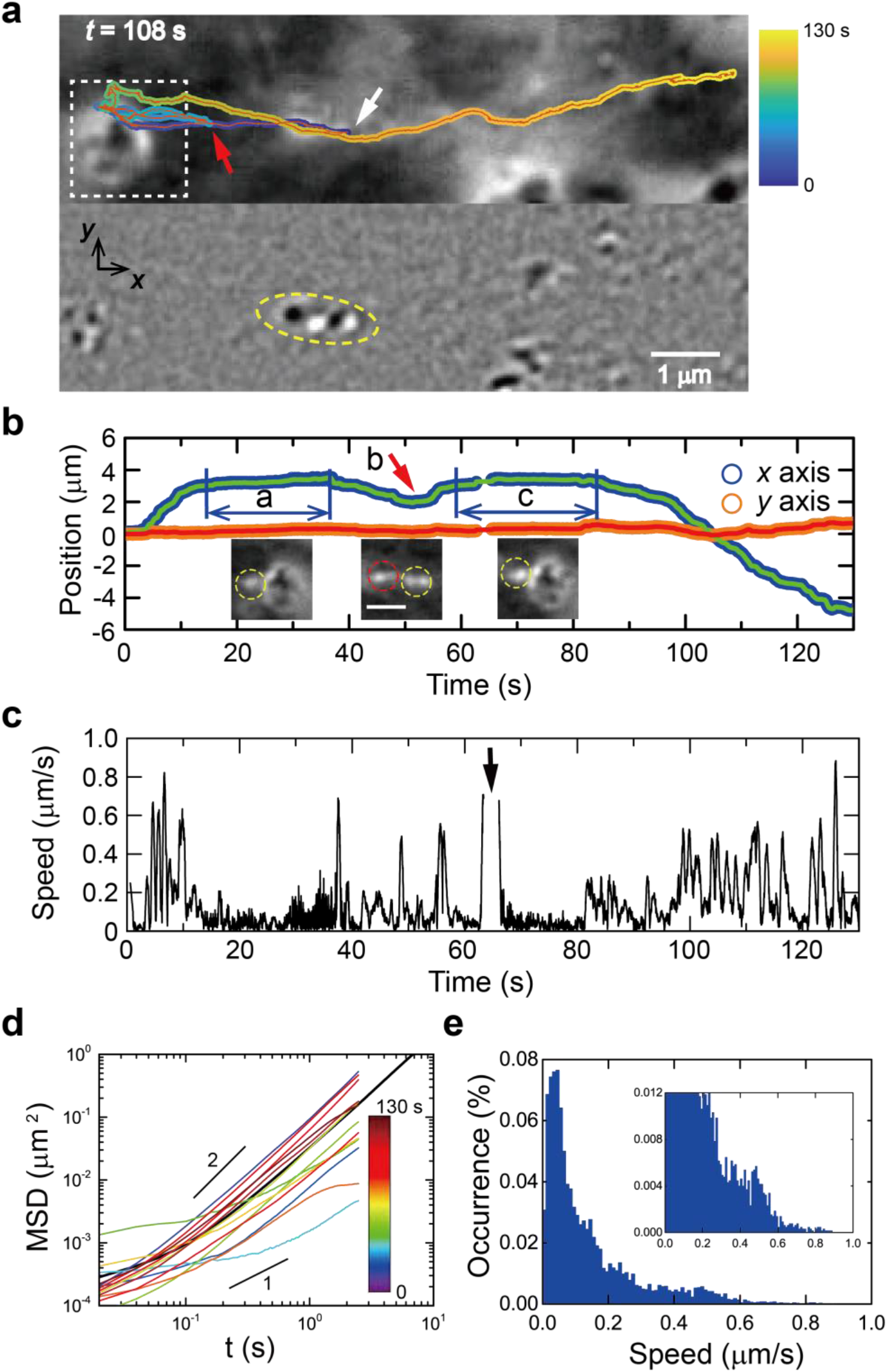
Dynamic events revealed by a directionally transported dimeric cargo. **a,** Trajectory of a dimeric cargo (yellow oval) showing a directed movement, turn-around, and intermittent pausing (upper: SBR-iSCAT snapshot, and lower: TD-iSCAT snapshot) (black-boxed area in Fig. 1d). Here, the white and red arrows indicate the starting point at t = 0 and turn-around point (time point ‘b’ in (b)) of the dimeric cargo, respectively. **b,** Raw and time-averaged positions of the dimeric cargo in x and y directions (x position: blue (raw) and turquoise (averaged); y position: orange (raw) and red (averaged)). During the pausing periods (‘a’ and ‘c’), the dimeric cargo (yellow circle in the left and right insets) interacts with a large spherical cytoplasmic structure placed on the right side. At the time point of ‘b’, the dimeric cargo (yellow circle in the middle inset) collided head-on with another cargo (red circle in the middle inset) moving in the opposite direction. **c,** Instantaneous speed of the dimeric cargo calculated from time-averaged data points (averaged over 50 raw data points for Δt = 1 s), which was measured with the interval of 0.02 s for the raw trajectory drawn in (a). When the dimeric cargo was located too close to the large spherical cytoplasmic structure, the automatic tracking program failed to work for a short period of time indicated by the arrow. **d,** MSD-vs.-time curves obtained from its whole trajectory (~ 130 s) drawn in (a) (black) and each time interval of 10 s (13 curves in different colors). **e,** Histogram of ‘instantaneous speed’ of the dimeric cargo obtained from (c). See also Extended Data Fig. 7, Supplementary Video 5-7.

Because the fluorescence-combined iSCAT microscopy captures the dynamic events of cargos regardless of f-PS labeling and detects both scattering from cargos and fluorescence from internalized f-PS beads simultaneously for precise specification, we could identify dimeric cargos straightforwardly (Extended Data Fig. 7a, Supplementary Video 8). A close inspection of the f-PS bead by iSCAT imaging revealed the f-PS bead as a part of a dimeric cargo. Interestingly, the dimeric cargo was put on hold because other cargos crossed the intersection ahead of it (Extended Data Fig. 7f). It was recently reported that intracellular cargos move together via ‘hitchhiking’. Some cargos literally can hitch a ride with nearby motile cargos already associated with molecular motors^35,36^. Intriguingly, iSCAT can directly visualize the outcome of hitchhiking, multimeric cargo, even when some cargos lack f-PS beads inside. A cargo trimer was also observed to move together along the same route (Extended Data Fig. 8, Supplementary Video 9). During translocation, one of the cargos was detached at *t* = 210.7 s, and the rest (dimer) continued to move and turned around at the intersection of the cytoskeleton from *t* = 215.7 s to 216.8 s. The hitchhiking process could be a rule rather than an exception to boost the efficiency of logistics by implementing specific adapter proteins bridging different vesicles^37,38^. Thus, it may be programmed for the cell’s benefit, which is rather analogous to carpooling.

Interestingly, the dynamic events described herein nicely match various occasions encountered during transportation: a car should detour at roadblocks, wait for other vehicles to cross at the intersection, and come to a stop at the end of the road. It is of note that a complete picture of cellular transportation is now made available by the brightfield iSCAT microscopy.

## Discussion

The fluorescence-based microscopic imaging provides direct evidence of various intracellular interactions among cargos, organelles, and cytoskeletons through multi-color labeling. Despite the strengths of target selectivity and high contrast, the fluorescence microscopy leaves the intracellular space surrounding labeled targets appearing dark, losing the information about physical interactions, which are potentially crucial for understanding the behavior of labeled targets. In contrast, we here demonstrated how iSCAT microscopy could effectively reveal critical information about intracellular cargo transport, which cannot be easily studied with fluorescence microscopy as well as ordinary bright-field microscopy. Instead of pursuing chemical contrast using fluorophores, iSCAT utilizes the dynamic feature of transported cargos to discriminate them from other cytoplasmic objects with similar shapes and iSCAT contrasts. One of the most notable advantages of iSCAT live-cell imaging is that it allows us to collect a large amount of data by detecting scattering signals from all kinds of cargos moving directionally along a cytoskeletal track over a long period of time. In the present study, we obtained about 10-million localization points from ~ 90,000 consecutive frames taken at 50 Hz (~ 30 min) on the same area (field of view: 20 × 20 μm^2^) of a single cell. The frame rate in the current iSCAT setup can be as high as ~ 1 kHz^28^, indicating that the same amount of data (~10-million localization points) can be acquired in 1.5 min. Even with the recent breakthrough in the imaging rate of fluorescence microscopy, up to a few hundreds of frames/s at diffraction-limited resolution^14^, it is hard to imagine that the state-of-the-art fluorescence microscopy can achieve the same level of imaging rate and a similar amount of image data from the same area of a single cell as presented in our iSCAT study.

The massive localization of cargos by iSCAT literally pictures the traffic of cargos and their underlying cytoskeletal highways. Although our cargo-localization iSCAT microscopy constructs the critical visual information with, in principle, the same point localization method generally used in super-resolution fluorescence microscopies^39–42^, it reveals quite different features for the following reasons. First, the current super-resolution fluorescence microscopy shows the whole meshwork of cytoskeletal material by accumulating localization points detected from photoswitchable dyes, directly labeled to major components of the cytoskeleton. In contrast, the cargo-localization-based iSCAT microscopy reconstructs the network from numerous localization points taken from actively transported cargos along the cytoskeletal passage, not from the cytoskeleton itself. Thus, it displays the cytoskeletal pipelines that are being actively used at the moment of observation, rather than the whole structure of a cytoskeletal network. Second, as a label-free, high-throughput imaging tool for extensive data acquisition, iSCAT captures the long-time evolution of cargo traffic flows on cytoskeletal highways. In contrast, present-day super-resolution fluorescence imaging is intrinsically limited due to the photobleaching of fluorophores. Recently, several nonlinear vibrational microscopies with chemical selectivity without fluorophore labeling have been introduced for live-cell imaging at a high spatial resolution^43–46^. However, these techniques are insufficient to capture the fast dynamics of cargo transportation in real-time. Thus, we believe that the functional imaging demonstrated here is unprecedented in fluorescence or vibrational microscopies.

In fact, these point-localization imaging methods use point-generation strategies with notably different underlying processes and features. In fluorescence-based single-molecule localization microscopy (SMLM), point generation is mediated by the stochastic activation of fluorophores and their subsequent emission, or the stochastic binding and unbinding of fluorophore-labeled probes^39,41,47,48^. The high overall density of points required for a detailed image and the sparsity of points at each frame necessary for reliable localization are simultaneously fulfilled by stochastic switching of target spots. In our cargo-localization iSCAT, point generation is mediated by spatial displacement of cargos in a manner somewhat similar to the (un)binding-based SMLM. The two motion-based localization strategies also exhibit notable differences: in the cargo-localization iSCAT, the displacement of targets occurs in the view field continuously and laterally and is mainly induced by the motor-driven active mechanism. Both diffusion-based passive and locomotion-based active mechanisms contribute to point localization and contrast, but likely in different time domains.

In one previous study that utilized an *in vitro* model system for motor protein transport, motor proteins traveling along microtubules were shown to undergo traffic jams as characterized by an abrupt increase in the density of motors and an associated abrupt decrease in motor speed^9,10^. Although this study provided some insights into traffic jams experienced by motor proteins at the molecular level, the *in vitro* models might not fully mimic the intrinsic complexity of the crowded cellular environment^49^. In contrast, our *in vivo* study showed that intracellular cargos regularly experience heavy traffic throughout the cytoskeletal network. For example, various dynamic events in cargo transport such as pausing, turn-around, and jamming are elucidated by obstacles that block the progress of the cargo, head-on collision with other cargos coming from the opposite side, or abrupt halt at the intersection of cytoskeletal highways. The physical association of cargo with the local cellular environment would be easily missed in fluorescence imaging. Surprisingly, a cell has inherent strategies for efficient cargo delivery to the destination through a heavily crowded cellular environment. In the short time scale, the traffic of cargos appears to be entangled and stagnant, but in the long timescale, they collectively move in the same direction as a cluster of cargos. In some cases, two or more cargos move together, seemingly being directly connected. Our experimental observations strongly support the recent hypothesis of intracellular transport by hitchhiking as one strategy to increase the overall transport rate^35,36^. As summarized above, the iSCAT approach would provide the unique opportunity to investigate the time evolution of nanoscopic cellular constituents and cytoplasmic context at high spatiotemporal resolution.

In this study, we identified the locations of intracellular cargos by using label-free iSCAT microscopy in conjunction with the image processing methods suitable for extracting the dynamic features of moving cargos from a complex cytoplasmic landscape. In general, the highly praised assets of the iSCAT technique, such as extreme sensitivity, concomitant fast-tracking capability, and quantitative measure of molecular mass, are significantly compromised when it is applied to the study of a living cell, which is highly inhomogeneous with numerous cytoplasmic objects with optical heterogeneity. The limitations of iSCAT, a lack of chemical specificity, preference for dynamic targets, and reduced performance in non-uniform environments, can be alleviated by integrating complementary imaging methodologies such as fluorescence microscopy, coherent Raman imaging, or IR photothermal microscopy.

## Methods

### Cell Culture procedures

COS-7 cells, monkey kidney fibroblast cell line, were cultured in a 35-mm confocal dish (SPL, Korea) pre-coated with 0.1 mg/ml poly-D-lysine (P7886, Sigma-Aldrich, USA) for good cell attachment. The seeding density of COS-7 cells was 3 × 10^5^ cells per dish. Then, cells were maintained at 37 °C in a humidified 5 % CO2 and 95 % air atmosphere in Dulbecco’s modified eagle medium (DMEM) (Gibco) supplemented with 10 % fetal bovine serum (FBS, Gibco) and 1 % penicillin/streptomycin (Gibco) for 1 day. Before iSCAT imaging, 20-nm f-PS beads were seeded into the sample dish and incubated for 6 hrs. It is of note that f-PS beads incubated with COS-7 cells are spontaneously internalized, trapped in vesicles, and then actively transported along cytoskeletal networks by motor proteins, as schematically depicted in Fig. 1b. Thus, directional cargos containing f-PS beads are likely intracellular vesicles. Then, the sample dish was loaded into a mini-incubating chamber (Chamlide, Live Cell Instrument, Korea) mounted on the piezo stage (MZS500, Thorlabs, USA) to maintain the temperature and pH in the sample dish during a long-term iSCAT live-cell imaging.

### iSCAT setup

The fluorescence-combined iSCAT microscopy setup is illustrated schematically (Extended Data Fig. 1). A continuous-wave diode laser (OBIS-FP-647LX, Coherent, USA) illuminates the sample area by raster scanning a weakly focused beam with a two-axis AOD (DTS-XY400-647, AA optoelectronics, France). The telecentric lenses (T1 and T2, *f* = 500 mm, AC254-500-A, Thorlabs, USA) are used to image the input beam onto the back focal plane of the high numerical aperture (NA) oil immersion objective lens (100× UPlanSApo, 1.4 NA, Olympus, Japan). The reflected and the scattered lights from the sample are collected by the same objective, reflected by the beam splitter (BS, BSW-20R, Thorlabs, USA), and imaged with the tube lens (TL1, *f* = 750 mm, AC254-750-A, Thorlabs, USA) onto an sCMOS camera (PCO.edge 4.2, PCO, Germany). The field of view is about 23 × 23 μm^2^. In order to detect fluorescent signals from f-PS beads, two dichroic mirrors (DM1 (FF495-Di03, Semrock) and DM2 (FF552-Di02, Semrock)) were introduced between the objective lens and the beam splitter. DM1 is used to reflect the excitation beam into the main beam path, and DM2 is used to reflect the fluorescent signal out of the beam path and into the detector. The light from an LED (SOLIS-3C, Thorlabs, USA) filtered by an optical bandpass filter (F1, FF02-482/18-25, Semrock, USA) is used to excite f-PS beads and the fluorescent signal from them, selected by DM2 and an optical bandpass filter (F2, FF02-520/28-25, Semrock, USA), is projected onto an EMCCD via a tube lens (TL2, *f* = 400 mm, AC254–400-A, Thorlabs, USA). The resulting field of view is about 32 × 32 μm^2^.

### SBR-iSCAT image processing

In principle, the iSCAT image is obtained by combining the scattering field from intracellular target objects with the reference field reflected at the interface between the glass and the culture medium. The intensity on the detector is given by

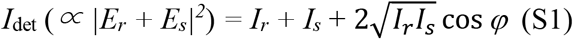

where *Er* and *E_s_* are the reference and scattered light fields, respectively, and *φ* is the relative phase between the two fields. As the scattering intensity (*I_s_*) is negligible for small objects, the third interferometric cross term becomes the leading contribution from scattering objects. The iSCAT contrast (*C*) is defined as

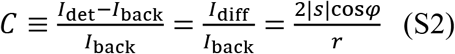

where *r* and *s* are the reflection and scattering coefficient, respectively.

The raw images from iSCAT inevitably include static non-uniform background caused by uneven illumination or spurious speckles due to back-reflections or defects on optical elements placed along the beam path, which deteriorates the quality of raw images. To eliminate such a time-invariant background noise, we employed the static background removed (SBR)-iSCAT image processing method (Extended Data Fig. 2a, c). First, a temporal-median cell image was generated by averaging 10 consecutive raw images (*Δt* = 0.2 s) taken at 50 Hz to reduce the level of shot noise fluctuations in iSCAT snapshots. Then, we obtained an SBR-iSCAT image by subtracting a temporal-median background image from the temporal-median cell image to obtain the differential image and dividing the differential image by the background image, as written in Eq. S2 above. The temporal-median background image was averaged for a stack of ~ 3,000 images obtained from a cell-free zone on a glass substrate. The quality of images is significantly improved thereby, and various cellular structures are clearly revealed in enhanced contrast by merely removing the stationary background with reduced shot noise.

### TD-iSCAT image processing

Even with the enhanced contrast in the SBR-iSCAT image, it is still difficult to identify intracellular cargos in the cytoplasmic space full of unidentified subcellular objects with similar shape and contrast. To capture dynamic cargos from the complex cytoplasmic landscape, we used the time-differential (TD)-iSCAT image processing method (Extended Data Fig. 2b, d) instead, highlighting dynamic cellular features by suppressing quasi-static iSCAT signals from relatively stationary cellular substance. First, we obtained a temporal-median cell image from 20 consecutive frames (*Δt* = 0.4 s) taken at 50 Hz. By dividing a temporal-median image by another such image taken after the time interval *(ΔT)* of 0.2 s, a TD-iSCAT image was obtained. In the image, individual cargos appear as a pair of dark and bright spots, indicating the early and late positions in the time interval averaged over for each image (*Δt* = 0.4 s). In contrast, other static or randomly moving cellular entities fade or disappear due to signal cancellation. In the showcase of a bi-directionally moving cargo, the dipolar look of a cargo conveys the direction of motion, indicated by the arrows, which is from dark to bright spot (Extended Data Fig. 2d). Compared to this, the corresponding SBR-iSCAT images fail to show the cargo due to irregular cytoplasmic background (Extended Data Fig. 2c). In the TD-iSCAT image sequence, it disappeared instantaneously at *t* = 1.96 s (Extended Data Fig. 2d) when the cargo changed the direction of motion, indicating that it intermittently stopped on a cytoskeleton track. The separation of the bright and dark spots corresponds to the displacement of the cargo occurring over *Δt*. Similar to a manufactured vehicle, the cargo slows down until it stops, turns around, and speeds up in the opposite direction (Extended Data Fig. 2d). For a backward motion, cellular cargos would have two choices: engaging motor proteins of opposite directionality on the same track or, less likely, taking another track of opposite polarity.

### Cargo localization and tracking via the MOSAIC ImageJ plugin

The detection and tracking of multiple cargos from iSCAT images was achieved by the MOSAIC ImageJ plugin^29^. Although this plugin was originally developed as an analytic tool for fluorescence microscopy images, we found that it is equally well suited for the analysis of cargos captured in both SBR- and TD-iSCAT images. There are three parameter values to be set for cargo identification: particle radius (*r*), cutoff score (*s_cut_*), and intensity percentile (*I_p_*). The value of particle radius was set to be the diffraction limit, 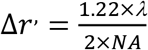, where the numerical aperture (NA) and *λ* used for iSCAT detection are 1.49 and 647 nm, respectively. Thus, *Δr′* is about 264 nm, corresponding to ~ 7 pixels in our iSCAT setup (1 pixel ~ 40 nm). Accordingly, larger cargos with a radius > 7 pixels were excluded, and two (or more) cargos less than 7 pixels apart could be counted as a single cargo in our analysis. The cutoff score for non-particle discrimination was not applied in our cargo identification and set to ‘0’. The intensity percentile is the parameter to determine which bright pixels are accepted as cargos. While the dynamic cellular features are only highlighted in TD-iSCAT cell images, SBR-iSCAT cell images reveal the whole cellular landscape, including static or randomly moving objects in enhanced iSCAT contrast. Thus, the intensity percentile was set to different values: *I_p_* = 5 and 0.3 for SBR- and TD-iSCAT images, respectively.

For cargo localization to reveal the underlying active cytoskeletal highways, the spatial coordinates of individual cargos are determined in the TD-iSCAT image with the following localization precision,

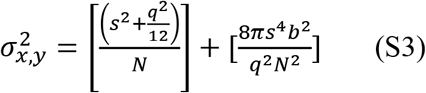

where s is the standard deviation of the point-spread function, N the total number of detected photons of the scattered light, q the equivalent dimension of a pixel in the sample space (~ 40 nm), and b the background noise per pixel^18,50,51^. Considering the full well capacity of the camera (~ 30,000 e^−^) given by the manufacturer’s datasheet, the values of N and b are assumed to be about 7000 and 1500, respectively, measured from a cell-free background area illuminated with light of 1 mW. Then, a pair of bright and dark spots revealed in a TD-iSCAT image were separately fitted with the following 2D Gaussian function,

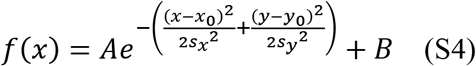

where *A* is the height of the peak with respect to the base level B, x0 and y0 are the positions of the center of the peak, and sx and sy the standard deviation of the position of a bright (or dark) spot (Extended Data Fig. 3). From our measurements, the value of SNR is ~ 6 (or 4) for a bright (or dark) spot, which allows localization of the center of the Gaussian peak with a precision of ~ 10 (or 15) nm.

### Instantaneous speed and MSD measurements from cargos’ trajectories

To obtain the trajectory from each individual f-PS bead-bearing cargo, the position of the f-PS bead was manually (Fig. 1f) or automatically (Fig. 1g) tracked from fluorescence images taken at 10 Hz. To automatically track the motion of an f-PS bead using the MOSAIC ImageJ plugin, we set *r* =7, *s_cut_* =0, and *I_p_* = 1. Trajectories from individual f-PS bead-free cargos (Figs. 3b, e, and 4a) were automatically obtained by applying the plugin to SBR-iSCAT images instead of TD-iSCAT images because the TD-iSCAT scheme fails to track cargos’ locations faithfully whenever they are intermittently paused. The instantaneous speeds (*v*) from the trajectories acquired in SBR-iSCAT images (Figs. 3g and 4c) were calculated from

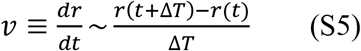

where *t* and *r*(*t*) are the time taken and the position of a particle at t, respectively. Here, *ΔT* is given by *1/f* (Hz), which is 0.02 s. However, the value of *v* calculated from a raw trajectory varies considerably, likely for the inaccuracy in finding the center position of cargo due to thermal noise. To suppress noise in speed measurement, we calculated the value of *v* from a 50-point time-averaged trajectory, the cargo position of which was given by the mean position of the cargo from 50 consecutive raw images (Extended Data Fig. 6). Markedly different from the raw and 10-point time-averaged trajectories, the 50-point time-averaged trajectories show that the baseline or minimum of *v*, which would be the value of speed at the moment of intermittent pause, is close to ‘0’.

The MSD at time t is defined by the ensemble average of the displacement squared over t:

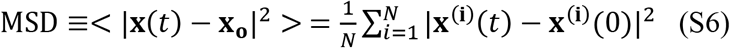

where *N* is the number of particles to be averaged, 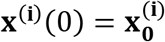 is the reference position of the i-th particle, and **x**^(1)^(*t*) is the position of the i-th particle at time t. A long-time particle tracking (> 130 s) necessary for MSD analysis is feasible for a single f-PS bead or an intracellular cargo as demonstrated with fluorescence imaging (Fig. 1g, h) or SBR-iSCAT time-lapse imaging (Fig. 4b, d). In contrast, cargo tracking is severely hampered in the bustling area where cargos frequently jam because the locations of cargos are frequently overlapped there. Thus, we selected a relatively short trajectory of each cargo for MSD analysis, continuously tracked for 150 consecutive frames (or 3 s) in the jammed area (Fig. 3b, c).

### Illustrations

All figures were created using OriginPro (Origin lab), Matlab (MathWorks), and Illustrator software (Adobe).

### Quantification and statistical analysis

The average speed of cargos moving on cytoskeletal highways is presented in the form of mean ± standard deviation and mentioned in the text. All trajectories of cargos were obtained from the same area (field of view: 20 × 20 μm^2^) of a single cell in Fig. 1c. From this area, ~ 10-million localization points automatically identified with the Mosaic ImageJ plugin were used to reconstruct the cytoskeleton network.

## Acknowledgments

This work was funded by IBS-R023-D1 (MC) and NRF-2019R1A2C1089808 (SH); Global Research & Development Center Program (2018K1A4A3A01064272, SH) through the NRF funded by the Ministry of Science and ICT.

## Author contributions

Conceptualization: JP, SH, MC. Methodology: JP, IL, MM. Investigation: JP, IL, MM, SH, MC. Writing – original draft: JP, SH, MC. Funding acquisition: SH, MC. Supervision: SH, MC.

## Additional information

**Supplementary information** The online version contains supplementary material

**Correspondence and requests for materials** should be addressed to S.-C. Hong or M. Cho.

